# Pupil size predicts the onset of exploration and changes in prefrontal dynamics

**DOI:** 10.1101/2023.05.24.541981

**Authors:** Akram Shourkeshti, Mojtaba Abbaszadeh, Gabriel Marrocco, Katarzyna Jurewicz, Tirin Moore, R. Becket Ebitz

## Abstract

In uncertain environments, intelligent decision-makers exploit actions that have been rewarding in the past, but also explore actions that could be even better. Several neuromodulatory systems are implicated in exploration, based, in part, on work linking exploration to pupil size–a peripheral correlate of neuromodulatory tone and index of arousal. However, pupil size could instead track variables that make exploration more likely, like volatility or reward, without directly predicting either exploration or its neural bases. Here, we simultaneously measured pupil size, exploration, and neural population activity in the prefrontal cortex while two rhesus macaques explored and exploited in a dynamic environment. We found that pupil size under constant luminance specifically predicted the onset of exploration, the first exploratory trial in a sequence, beyond what could be explained by reward history. Pupil size also predicted disorganized patterns of prefrontal neural activity at both the single neuron and population levels, even within periods of exploitation. Ultimately, our results support a model in which pupil-linked mechanisms promote the onset of exploration via driving the prefrontal cortex through a critical tipping point where prefrontal control dynamics become disorganized and exploratory decisions are possible.

**Significance Statement:** Humans and other animals learn about the world through exploration: through making decisions that offer the opportunity to learn and discover, even when these decisions are not the best option in the moment. Neuroscience research has historically focused on understanding good choices, delivering many key insights into the neural mechanisms involved in these calculations. However, much less is known about how the brain generates exploratory decisions. This study identifies certain “early warning signs” of exploratory decisions in the brain and body, including certain signals in size of the pupil and the speed of neural activity in the prefrontal cortex. These early warning signs suggest that exploration may be the result of a critical tipping point in prefrontal brain states.

## Introduction

Many decisions maximize immediate rewards. However, in uncertain or changing environments, it is important to sacrifice some immediate rewards in order to learn about the value of alternative options and discover new, more valuable strategies for interacting with the world. In short, in complex environments, intelligent decision-makers exploit rewarding strategies, but also explore alternative strategies that could be even better.

Because exploitation maximizes immediate reward, it can rely on the same value-based decision-making processes that have been the subject of neurobiological studies for decades (Ding and Hikosaka, 2006; Jurewicz et al., 2022; Platt and Glimcher, 1999; Roesch and Olson, 2007; Schultz et al., 2008). However, we are only just beginning to understand the neural bases of exploration (Costa and Averbeck, 2020; Daw et al., 2006; Pearson et al., 2009; Wilson et al., 2021, 2014). One clue is that many organisms seem to explore via random sampling (Gershman, 2019; Wilson et al., 2021, 2014). Randomness is a critical component of exploratory discovery in bird song and motor learning (Fiete et al., 2007; Wu et al., 2014), it can perform about as well as more sophisticated strategies in many environments (Dayan and Daw, 2008), and humans and other primates tend to explore randomly even when more sophisticated strategies are available (Ebitz et al., 2018; Wilson et al., 2014). There is some neurobiological evidence linking random exploration to disorganized activity patterns in the prefrontal cortex (Ebitz et al., 2019, 2018; Muller et al., 2019; Wilson et al., 2021), but we still do not understand the proximate causes of random exploration and its neural correlates in the prefrontal cortex.

One promising hypothesis is that exploration could be under the control of some process(es) linked to pupil size. Pupil size under constant luminance is a peripheral index of autonomic arousal (Bradley et al., 2008; Ebitz and Moore, 2019; Loewenfeld, 1999) that also predicts widespread changes in neural population activity (McGinley et al., 2015; Reimer et al., 2014)– including in regions implicated in decision-making noise (Ebitz and Platt, 2015; Tervo et al., 2014). Among other neuromodulators (Gilzenrat et al., 2010; Koss, 1986; Reimer et al., 2016), pupil size is correlated with central norepinephrine (Costa and Rudebeck, 2016; Joshi et al., 2016): a catecholamine that flattens neuronal tuning functions (Martins and Froemke, 2015) and predicts “resets” in cortical networks (Aston-Jones and Cohen, 2005; Bouret and Sara, 2005). Behaviorally, pupil size predicts decision-making noise (Aston-Jones and Cohen, 2005; Ebitz et al., 2014; Eldar et al., 2013; Gilzenrat et al., 2010; O’Reilly et al., 2013; Wilson et al., 2021), especially errors of reward-maximization (Jepma and Nieuwenhuis, 2011a) and task performance (Ebitz et al., 2014; Ebitz and Platt, 2015). Some of these “errors” may be caused by exploratory processes (Ebitz et al., 2019; Jepma and Nieuwenhuis, 2011a; Pisupati et al., 2021).

There is a plausible alternative interpretation of this data: perhaps pupil size only tracks the variables that make exploration more likely. Pupil size under constant luminance increases with the volatility of reward environments, the surprise of reward outcomes, novelty, uncertainty, and context changes (Clewett et al., 2020; Filipowicz et al., 2020; Graves et al., 2021; Preuschoff et al., 2011; Slooten et al., 2018; Yokoi and Weiler, 2022): all variables that make exploration more likely. However, it is often unclear whether the pupil is tracking these variables or instead directly predicting behavioral changes like increased learning, decision-noise or exploration (Nassar et al., 2012; O’Reilly et al., 2013; Urai et al., 2017). Fortunately, recent results suggest that at least some exploration appears to occur tonically, regardless of these variables (Ebitz et al., 2019; Pisupati et al., 2021; Wilson et al., 2021). Further, in parallel, new computational approaches allows us to determine when exploration is occurring independently of the reward-based computations thought to drive it (Chen et al., 2021; Ebitz et al., 2020, 2019, 2018). This means that it is now possible to determine whether pupil size predicts exploration itself or instead simply tracks the variables that make exploration more likely.

Here, we measured pupil size and recorded from populations of prefrontal neurons while two rhesus macaques performed a task that encouraged exploration and exploitation. We found that pupil size under constant luminance was larger during explore choices than exploit choices. However, the temporal relationship between pupil size and exploration was both precise and complex: spontaneous oscillations in pupil size entrained the onset of exploration. Together, these results support the hypothesis that pupil-linked processes drive the prefrontal cortex through a critical tipping point that permits exploratory decisions.

## Results

Two male rhesus macaques performed a total of 28 sessions of a classic explore/exploit task: a restless three-armed bandit (subject B: 10 sessions, subject O: 18 sessions; a total of 21,793 trials). Some analyses of this dataset have been reported previously (Ebitz et al., 2018), but the pupil data has not been analyzed previously and all analyses presented here are new. In this task, the reward probability (value) of three targets walks randomly and independently over time (**Figure 1A**). This means that the subjects have to take advantage of valuable options when they are available (exploit), but also occasionally sample alternative options to determine if they have become more valuable (explore).

**Figure 1.**
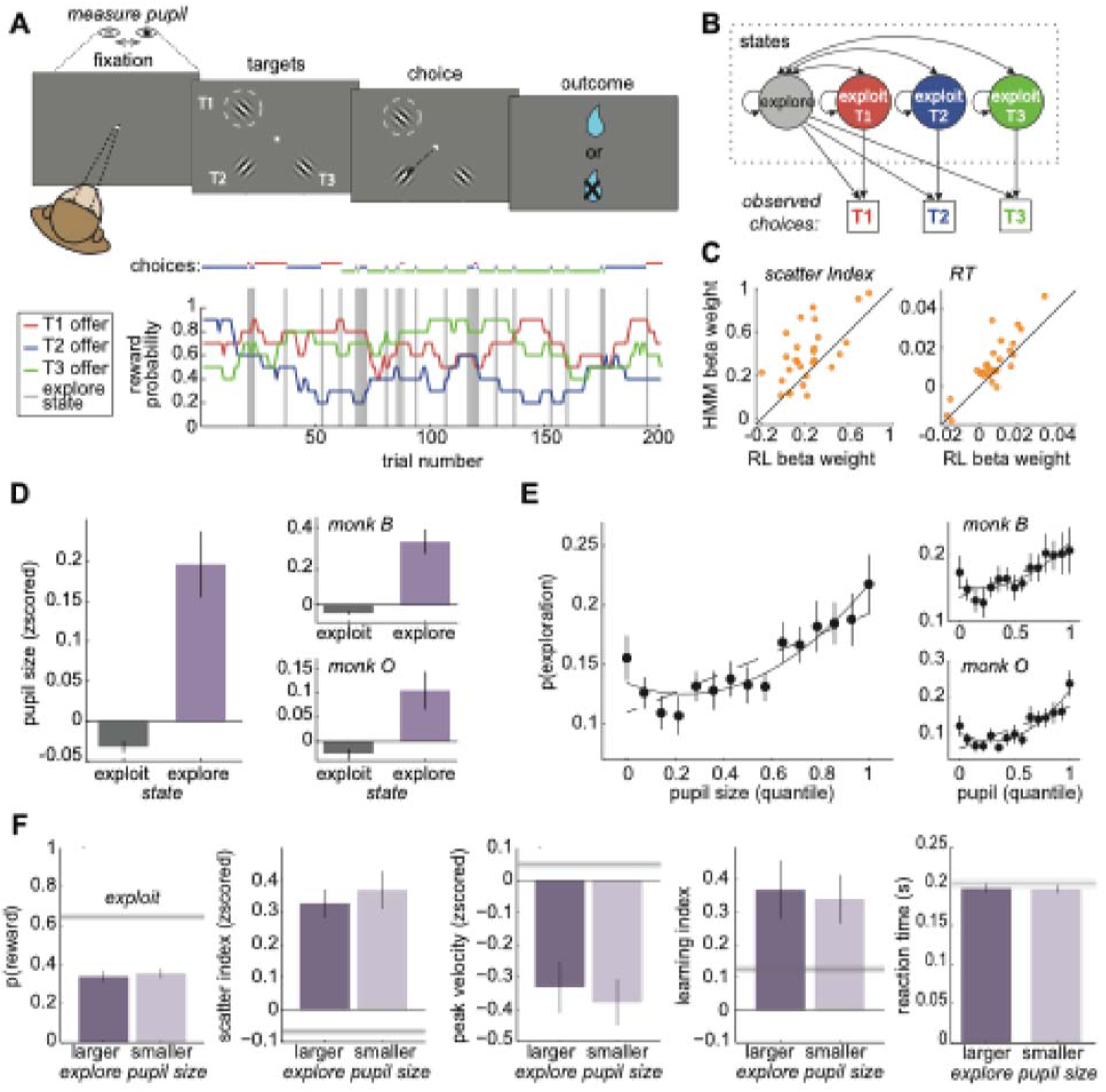
Task design and pupil. A) Top: Subjects made saccadic choices between three identical options (T1, T2, and T3). One of the options (e.g., T1 in this example trial) was located in the receptive field of a neuron in the frontal eye field (FEF; dotted circle). Bottom: Reward probabilities for the 3 options (lines), with choices overlaid (dots) for 200 example trials. Gray bars = explore-labels. B) The HMM models exploration and exploitation as latent goal states underlying choice sequences. C) Comparison of regression coefficients for HMM-inferred and RL-inferred explore choices, predicting either the disorganization of neural population responses (“scatter index”; see Methods; Ebitz et al., 2018) or response time. Beta weights were obtained from session-level generalized linear models (GLMs) with explore state as the predictor (0 = exploit, 1 = explore). Separate GLMs were fitted using explore labels from the HMM and from an RL-based model. D) Average pupil size on explore and exploit choices. Right: Same for individual subjects. E) The probability of explore choices as a function of pupil size quantile. Dotted line: linear GLM fit. Solid line: quadratic fit. Right: Same for individual subjects. F) Several behavior measures compared across median-split large- and small-pupil-size explore choices. Left to right: reward probability, a one-trial-back learning index (see Methods), saccadic peak velocity of saccades, the scatter index, and reaction time. No significant differences between pupil bins. The grey line is the mean ± SEM for exploit choices. Error bars depict ± SEM throughout.

Rather than instructing subjects to explore and exploit, this task takes advantage of the subjects’ natural tendency to alternate between exploration and exploitation in a changing environment. We have previously shown that both monkeys and mice exhibit 2 behavioral modes in this task: one exploitative mode in which they repeatedly choose the same option— learning little but maximizing reward—and one exploratory mode in which they alternate rapidly between the options—choosing randomly with respect to rewards and learning rapidly (Chen et al., 2021; Ebitz et al., 2018). We infer which of these modes is driving behavior with a hidden Markov model (HMM; **Figure 1B**; see **Methods**). This approach models the exploratory and exploitative modes as latent goal states and the maximum *a posteriori* goal is taken as the state label for each choice. We have previously shown that this method identifies explore/exploit state labels that match normative definitions (Chen et al., 2021; Ebitz et al., 2018) and explain variance in prefrontal neural activity that cannot be explained by reward value, reward history, and switch/stay decisions (Ebitz et al., 2018). This task design naturally elicits exploration and exploitation, allowing us to investigate variability in pupil size and neural activity under both conditions.

Some previous studies used a different method to identify exploratory choices (Daw et al., 2006; Jepma and Nieuwenhuis, 2011a; Pearson et al., 2009). These studies fit a reinforcement learning (RL) model to the behavior and identified the choices that are not consistent with the model’s subjective values as exploratory. However, this previous RL-based approach (1) equates exploration with errors of reward maximization, not a goal that is orthogonal to reward maximization, and (2) its accuracy depends on precise knowledge of the computations involved in the choice, which are highly variable, both across individuals and over time (Chen et al., 2021, 2021; Kaske et al., 2022). The HMM approach, conversely, makes no assumptions about the computations involved in the choice and identifies choices that are orthogonal to reward value, not anti-correlated with it (Chen et al., 2021; Ebitz et al., 2018). Here, we found that state labels from the HMM method explained more variance in behavior and neural activity than choice labels from the previous, RL method (**Figure 1C**; response time: both subjects, paired t-test: p < 0.005, t(27) = 3.41, the mean difference of beta weights = 0.004, 95% CI = 0.002 to 0.007; scatter index [(Ebitz et al., 2018)]: both subjects, paired t-test: p < 0.001, t(27) = 3.84, the mean difference of beta weights = 0.15, 95% CI = 0.07 to 0.24: see **Methods**). In short, we find that the HMM approach is a more robust and accurate method, with better face validity, than the RL-based method for identifying explore choices. Therefore, here, we used this more precise approach to determine whether physiological signals, like pupil size, reliably track exploratory behavior.

### Pupil size is larger during exploratory states

Having established that the HMM reliably distinguishes explore and exploit states, we next asked whether pupil size changes across these behavioral states. Previous work using RL-based labels reported that pupil size under constant luminance is larger during exploration than exploitation (Jepma and Nieuwenhuis, 2011b). We therefore tested whether this pattern holds using HMM-based labels in our dataset. Indeed, we found that pupil size at fixation (see **Methods**) was larger on explore-labeled trials than exploit-labeled trials in both subjects (**Figure 1D**; both subjects, paired t-test: p < 0.0001, t(27) = 4.95, mean offset = 0.23, 95% CI = 0.13 to 0.32; subject B: p < 0.001, t(9) = 5.50, mean offset = 0.4, 95% CI = 0.24 to 0.57; subject O: p < 0.02, t(17) = 2.85, mean offset = 0.13, 95% CI = 0.03 to 0.23). Thus, pupil size was larger during exploratory choices identified with the HMM method.

However, the probability of exploration did not increase linearly as a function of pupil size (**Figure 1E**). A linear, first-order GLM confirmed that larger pupil size generally predicted more explore choices (both subjects: β = 0.084, *p* < 0.0001, AIC = –1053.70, n = 28 sessions). This relationship held when analyzed separately in each subject (subject B: β = 0.063, *p* = 0.002, AIC = –368.01, n = 10 sessions; subject O: β = 0.110, *p* < 0.0001, AIC = –249.46, n = 18 sessions). Yet the relationship was clearly nonlinear. A quadratic model provided a significantly better fit than the linear model for the combined dataset (2nd order GLM: β_₁_ = –0.081, *p* = 0.101; β_₂_ = 0.166, *p* = 0.0006; AIC = –1063.62, AIC weight for the quadratic model = 0.993), consistent with a U-shaped relationship. This U-shape was especially prominent in subject O (β_₁_ = –0.13, *p* = 0.090; β_₂_ = 0.240, *p* = 0.001; AIC = –258.01), although the quadratic model was not an improvement over the linear model in subject B (β_₁_ = –0.027, *p* = 0.709; β_₂_ = -0.091, *p* = 0.205; AIC = –367.63).

In order to determine whether this pattern was also apparent in the raw data (i.e. not the HMM-model labels), we next examined how pupil size predicted switching behavior—i.e., choosing a different option than in the previous trial. We again observed a U-shaped relationship between pupil size and the probability of making a switch choice in both subjects (1st-order GLM: β = 0.084, p < 0.0001, n = 28 sessions; subject B: β = 0.0800, p < 0.0001; subject O: β = 0.0807, p < 0.0001). A 2nd-order quadratic model provided a superior fit in both animals (both subjects: β_₁_ = –0.099, p = 0.023; β_₂_ = 0.184, p < 0.0001; subject B: β_₁_ = –0.069, p = 0.284, β_₂_ = 0.149, p = 0.010; subject O: β_₁_ = –0.093, p = 0.103, β_₂_ = 0.174, p = 0.001). Model comparison strongly favored the quadratic model (linear AIC = –1195.05; quadratic AIC = –1211.95; AIC weight for quadratic model = 0.9998). Thus, although pupil size tended to be larger during exploration than exploitation, its relationship with both exploration and switching was clearly U-shaped.

One possible explanation for the U-shaped pattern is that some “explore” choices—particularly those with small pupil size—reflect disengagement at low levels of arousal, rather than true exploration. However, if this were the case, then the valid, large-pupil explore choices would systematically differ from the false, small-pupil “explore” choices. They did not. To evaluate this possibility, we compared small- and large-pupil explore trials across several behavioral dimensions known to be sensitive to lapses in task engagement. These included reward rate, saccade velocity, the neural scatter index, a trial-wise learning index, reaction time—each previously associated with arousal, motivation, or task-related updating (Chen et al., 2021; Ebitz et al., 2018; Laurie et al., 2025). Small- and large-pupil explore choices (median split) were indistinguishable along several of the key dimensions that differentiate explore choices from exploit choices (**Figure 1F**). For example, both were equally likely to be rewarded (mean difference = 0.03 ± 0.24 STD) between large- and small pupil-explore choices (p > 0.4, t(1,27) = 0.75, paired t-test; AUC for discriminating explore and exploit = 0.65 ± 0.05 STD across sessions). Both had similar peak saccadic velocities (mean difference = -0.05 ± 0.23 STD, p > 0.2, t(27) = -1.08; explore/exploit AUC = 0.61 ± 0.10 STD) and both had more variability in neural population choice information (“scatter index”, mean difference = 0.03 ± 0.33 STD, p > 0.6, t(27) = 0.45; explore/exploit AUC = 0.60 ± 0.07 STD). Both had similar levels of reward learning (see **Methods**; the mean difference = -0.03 ± 0.57 STD, p > 0.7, t(27) = 0.27): in both cases, learning was substantially enhanced relative to the exploit choices (small-pupil, the mean difference from exploit = 0.24 ± 0.48 STD, p < 0.02, t(27) = 2.69; large-pupil, the mean difference from exploit = 0.21 ± 0.39 STD, p < 0.01, t(27) = 2.91). Reaction times were also similar across small- and large-pupil explore choices (mean difference = 0.01 ± 0.02 STD, *p* > 0.6, *t*(27) = 1.84; explore/exploit AUC = 0.58 ± 0.06 STD). These results are incompatible with the idea that either type of explore choice reflects disengagement in the task or that small- and large-pupil explore choices have different causes. Instead, we will see that the U-shape was due to the complex temporal relationship between pupil size and exploration.

### Pupil size, but not other measures ramp up before exploration

Pupil size ramped up across trials before exploration began in both subjects. After exploration, it shrank to below-baseline levels when exploitation resumed (**Figure 2A**). Here, “baseline” refers to a z-scored value of 0, computed by subtracting the session mean and dividing by the session standard deviation of pupil size (see **Methods**).This ramping meant that pupil size was larger not just during exploration, but also during the exploit choices immediately before exploration (both subjects, GLM slope = 0.01, p < 0.005, n = 28; subject B: beta = 0.02, p < 0.02, n = 10; subject O: beta = 0.01, p < 0.05, n = 18; average pupil size compared to the exploit choices, post-hoc paired t-tests, 1 trial before exploration mean = 0.12, p < 0.005, t(27) = 3.42; 2 trials mean = 0.09, p < 0.03, t(27) = 2.41; 3 trials mean = 0.03, p > 0.1, t(27) = 1.42; 4 trials mean = 0.05, p < 0.05, t(27) = 2.07). By the first exploit choice after exploration, pupil size had already begun shrinking to below-baseline levels (post-hoc paired t-tests, 1 trial after exploration mean = 0.03, p = 0.09, t(27) = 1.73; 2 trials after mean = -0.11, p < 0.02, t(27) = -2.67; 3 trials after mean = -0.16, p < 0.02, t(27) = -2.72; 4 trials after mean = -0.08, p > 0.2, t(27) = -1.27; 5 trials after mean = -0.16, p < 0.001, t(27) = -3.94; p-values are significant with a Holm-Bonferroni correction). The shrinking to below-baseline levels could suggest a refractory mechanism that would prevent exploration from re-occurring immediately after it happened.

**Figure 2.**
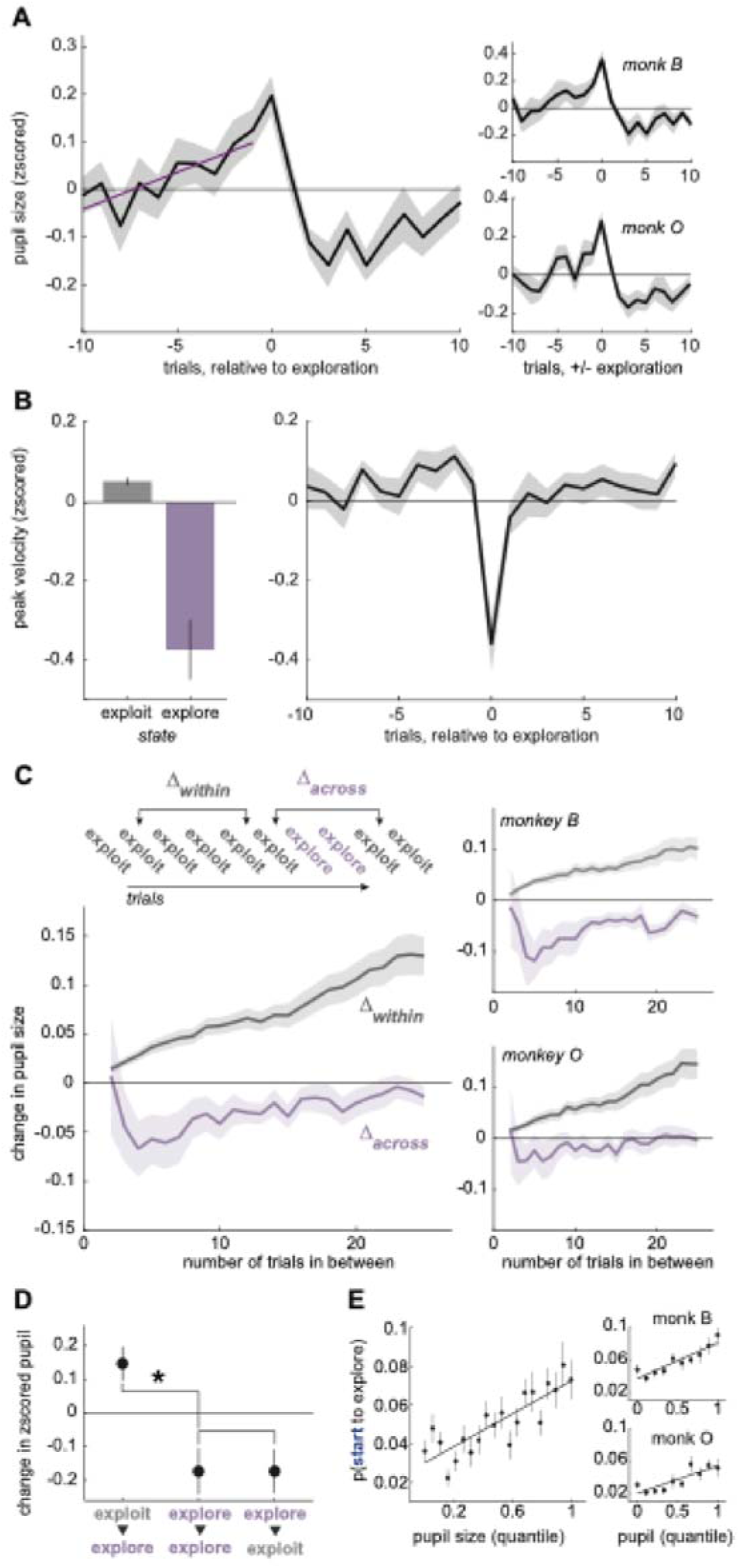
Pupil size ramps up before exploration and shrinks down after. A) Average pupil size for 10 trials before and 10 trials after explore choices. Purple line: GLM fit. Right: Same for each subject separately. B) Same as Figures 1D and 2A but for peak velocity rather than pupil size. C) Change in pupil size between exploit trials that are either in a single bout of exploitation (gray) or separated by explore trials (purple). Right: Same for each subject separately. D) Change in pupil size over certain pairs of trials: starting (exploit to explore), during (explore to explore), and leaving (explore to exploit) exploration. *p < 0.001 E) The probability of starting to explore as a function of pupil size quantile. Solid line: Linear GLM fit. Error bars and shaded regions depict mean ± SEM. Insets: Same analysis shown separately for each monkey.

To rule out potential confounds, we tested whether the pupil ramping and shrinking effects could be explained by misaligned labels or unrelated behavioral dynamics. We saw no evidence of ramping in peak saccadic velocity, another behavioral measure that differentiated explore trials and exploit trials (**Figure 2B**; no significant decrease from baseline 1 trial before, paired t-test: p > 0.7, t(27) = 0.42; a GLM was nonsignificant with the trend pointing in the opposite direction: 10 trials preceding exploration, beta = 0.008, p > 0.1) and no significant change from baseline afterward (not greater than the baseline during the 5 trials after exploration, when pupil shrinking was maximal, mean = -0.47 ± 0.15 STD, p > 0.9, one-sided t(27) = -1.67). We previously reported similar results for decoded choice probability and the scatter index (see **Methods**) in neurons of the frontal eye field (FEF), a prefrontal cortex region involved in directing gaze and attention (Ebitz et al., 2018). Thus, while pupil size ramped before exploration began and shrank afterward, the same was not true of other behavioral and neural variables, suggesting that these dynamics were not some artifact of misalignment.

### Pupil size generally ramps across trials, but resets with exploration

To better understand whether pupil ramping was a general feature of arousal dynamics or specific to exploration, we examined how pupil size evolved across trials with and without exploratory transitions (see **Methods**). When the subjects did not explore the pupil size increased steadily across trials (**Figure 2C**; both subjects, GLM: beta = 0.004, p < 0.0001; subject B: beta = 0.003, p < 0.0001; subject O: beta = 0.005, p < 0.0001, n = 25 lags over 28 sessions). This implies that the ramping in pupil size before explore choices may be a general dynamic of how pupil size evolves in the absence of exploration. However, a different pattern emerged when we looked at how the pupil changed between exploit trials that were separated by exploration. When two exploit trials were separated by at least one explore choice, pupil size was smaller on the second exploit trial (both subjects, GLM: beta = -0.09, p < 0.0001; subject B: beta = -0.14, p < 0.0001; subject O: beta = -0.07, p < 0.0001). Critically, passing through exploration only produced a baseline decrease in pupil size but did not alter the rate at which pupil size grew over trials (no significant interaction between slope and condition in both subjects, GLM: beta < -0.0001, p > 0.9; subject B: beta = 0.003, p < 0.05; subject O: beta = -0.002, p > 0.1; also nonsignificant on trials 5-25: both subjects: beta < 0.0005, p > 0.5). Therefore, pupil size tended to ramp across trials but exploratory choices temporarily decreased pupil size without disrupting this ramping in the long term.

### Pupil size specifically predicts the onset—not the maintenance—of exploration

Pupil size tends to be smaller after exploration, but this shrinkage could either be driven by the end of exploration (i.e. the start of exploitation) or it could begin shortly after the beginning of exploration itself. If the pupil starts to shrink only after exploration ends, it would support models that suggest that pupil size decreases with commitment to a new option or belief state (O’Reilly et al., 2013). Conversely, if the pupil shrinks immediately after exploration begins, it might suggest that pupil-linked mechanisms are important for initiating exploration, but not sustaining it. Our results were consistent with the latter hypothesis: the pupil immediately began shrinking as soon as exploration began, not when it ended (**Figure 2D**; mean change in pupil size between neighboring explore choices = -0.17, t-test, t(27) = -2.69, p < 0.02; 95% CI = -0.30 to -0.04). This was essentially identical to the magnitude with which the pupil shrank on exploit trials that followed explore trials (mean change = -0.17, t-test, t(27) = -2.56, p < 0.02; 95% CI = -0.31 to -0.03). Validating the ramping we observed with other methods, we also found that pupil size tended to grow on explore trials that followed exploit trials here (mean change = 0.14, t-test, t(27) = 2.96, p < 0.01; 95% CI = 0.04 to 0.24). Together, these results suggest that pupil size and pupil-linked mechanisms specifically predict the “onset” of exploration—the first exploratory trial in a sequence—and may not be important for sustaining exploration after the first explore choice.

The tendency of the pupil to shrink after the onset of exploration could explain the previously noted U-shaped relationship between pupil size and exploration. In this view, the small-pupil-size explore choices would be the later explore choices in a sequence and the larger pupil size explore choices would tend to be the first explore choice(s). Indeed, pupil size had a primarily linear relationship with the onset of exploration in both subjects (**Figure 2E**; 1st order GLM: β = 0.042, p < 0.0001, AIC = –1973.57). Adding a quadratic term did not substantially improve the model fit (β_₂_ = 0.038, p = 0.073; quadratic model AIC = –1974.81; ΔAIC = –1.24; AIC weight of quadratic model = 0.65; see **Methods**). This linear relationship was also observed in both monkeys individually (see **Figure 2E**, right panels). For subject B, a first order GLM confirmed a significant positive association (β = 0.044, *p* < 0.0001, AIC = –699.13), and adding a quadratic term did not improve the model fit (β_₂_ = 0.049, *p* = 0.0878; ΔAIC = –0.96). Similarly, for subject O, pupil size showed a significant linear relationship with exploration onset (β = 0.034, *p* < 0.0001, AIC = –460.29), and the quadratic model again provided no additional explanatory power (β_₂_ = 0.022, *p* = 0.402; ΔAIC = +1.30). These results confirm that the linear relationship between pupil size and the onset of exploration was robust across both subjects and not driven by outliers or subject-specific variability. Conversely, there was no special relationship between pupil size and probability of starting to exploit (1st order GLM: beta = 0.05, p > 0.05). Thus, pupil size specifically predicted the onset of exploration, rather than explore choices or state switches more generally.

If the U-shaped relationship between pupil size and exploration (**Figure 1E**) were driven primarily by later explore trials, it should remain evident after excluding onset trials from the analysis. Moreover, removing onsets should substantially reduce the slope of the linear effect. To test this, we repeated the analysis using only later explore trials. As expected, the linear slope decreased in both subjects (**Figure S1**): for subject B, β_1_ dropped from 0.063 (all explore trials) to 0.018 (excluding onsets), and for subject O, from 0.110 to 0.075. In contrast, the quadratic terms remained relatively stable: for subject B, β_2_ was 0.091 for all explore trials and 0.041 with onsets excluded; for subject O, β_2_ was 0.240 and 0.215, respectively. Importantly, the U-shaped relationship persisted when data were combined across both subjects (**Figure S1**), with the quadratic model significantly outperforming the linear model (β2=0.121,p=0.012\beta_2 = 0.121, p = 0.012β2=0.121,p=0.012; AIC = –878.78 vs. –874.55; AIC weight for the quadratic model = 0.893). These findings confirm that the nonlinearity observed in the original analysis (**Figure 1E**) was driven by the decrease in pupil size in later exploratory trials, whereas the onset of exploration had a largely linear relationship with pupil size.

### Exploration is gated by pupil-linked arousal, not just reward history

Although pupil size predicted the onset of exploration, it remained possible that this relationship was driven by a shared sensitivity to recent reward outcomes, since both exploration (Daw et al., 2006; Ebitz et al., 2018; Wilson et al., 2014) and pupil dilation (Bijleveld et al., 2009; Jepma and Nieuwenhuis, 2011b) tend to increase following reward omission. To determine if there was a direct effect of pupil-related processes on exploration, we compared pupil size across exploit trials before exploration with pupil size from matched trial sequences where exploration did not happen (see **Methods**). There was a significant increase in pupil size during the trials before exploration compared to “matched rewards” control trials (**Figure 3B**; GLM, beta = 0.025, p < 0.01, n = 28), suggesting that pupil size predicted the onset of exploration beyond what could be explained by reward. Again, pupil size ramped up over time (GLM, beta = 0.119, p < 0.02, n = 28), but this ramping did not differ between the traces (GLM, beta = 0.007, p > 0.5, n = 28). This implies that either reward history or time (i.e., the number of trials) may explain the pupil ramping before exploration, although there is still an offset in pupil size that predicts the onset of exploration above and beyond the effect of reward history.

**Figure 3.**
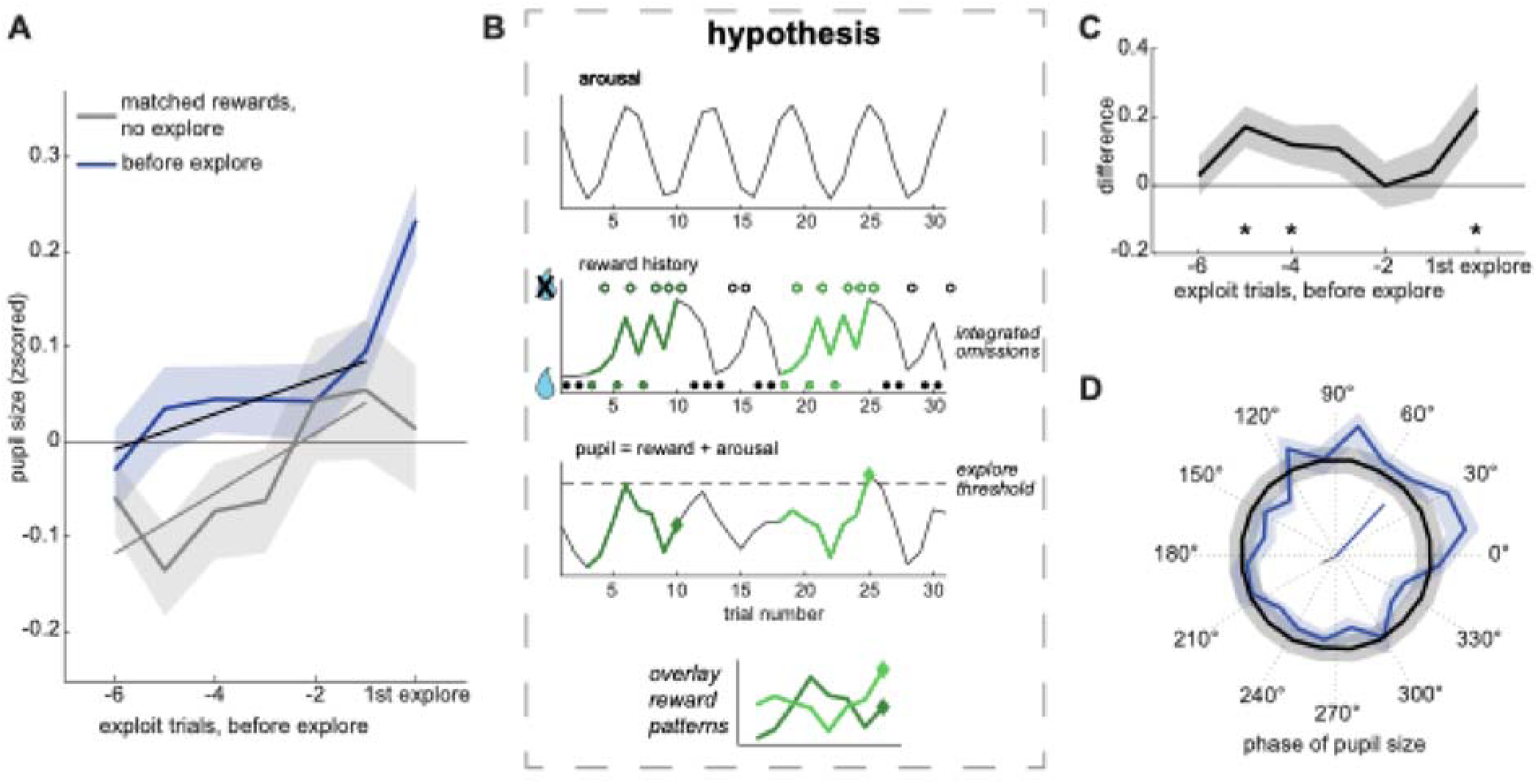
The onset of exploration is phase-locked to pupil size. A) Average pupil size over sequences of exploit trials before the onset of exploration (black line) and sequences with matched rewards, but no exploration at the end (gray line). Lines: GLM fit. B) Cartoon illustrating how oscillations in arousal (top) could interact with reward history (middle), to regulate exploration. The bottom panel illustrates a hypothetical pupil trace that has an additive effect of reward omissions and by oscillating arousal. Exploration (diamond shapes) begins when pupil size reaches a threshold (dotted line). Note that identical patterns of reward delivery and omission have different outcomes, depending on how they align with the phase of arousal (gray = no exploration, blue = exploration). C) Difference in pupil size between the traces in A. D) Phase distribution of pupil size at the onset of exploration (blue) and bootstrapped null distribution (black). The vectors at the center indicate the mean vector direction and length for the trials before exploration (blue) and the matched reward trials (gray). Shaded areas ± SEM throughout.

Visual inspection of **Figure 3A** suggested that there may be a phase difference in pupil size between trials where exploration began and matched reward trials where exploration did not begin. This led us to develop a novel hypothesis (**Figure 3B**): that rewards may interact with ongoing oscillations in pupil size. Due to delays in communication between the baroreceptor reflect and changes in heart rate, the sympathetic nervous system (Borjon et al., 2016; Japundzic et al., 1990; Julien, 2020, 2006; Kamiya et al., 2005; Liao et al., 2018) has a natural oscillation known as the Mayer wave, with a period of approximately 0.05–0.1 Hz (Borjon et al., 2016b; Julien, 2006). Critically, transitions in other behavioral states can be entrained by this oscillation, with eliciting stimuli causing transitions only at certain phases of arousal. This view predicts that the previous trials most predictive of the onset of exploration may actually be several trials prior to onset itself—during the periods in which the signals are most out of phase. Indeed, the trials in which pupil size best predicted the onset of exploration were not those immediately preceding it (e.g., trial t–1 or t–2), but rather trials t–4 and t–5 (**Figure 3C**; trial t–4, mean difference = 0.117, p < 0.05, t(27) = 2.09; trial t–5 = 0.170, p < 0.01, t(27) = 2.84).

The view that omitted rewards only evoke exploration when they coincide with particular phases of sympathetic arousal also implies that that the onset of exploration should be phase-locked to the Mayer wave frequency (see **Methods**). The median trial duration was ∼3 seconds (range = [2.2, 3.2]), so a ∼5-trial cycle would correspond to a 0.06–0.09 Hz oscillation, which aligns with the frequency range of the Mayer wave. We found that pupil phase at the onset of exploration was concentrated at the rising phase (**Figure 3D**; mean phase = 47.18°, Hodges-Ajne test, p < 0.01; vector length = 0.075; null = 0.026, 95% CI = 0.004 to 0.057, p < 0.0001). In contrast, pupil phases during reward-matched trials pointed in the opposite direction (mean phase = 207.58°; significantly different from onsets, p < 0.02, Watson’s U² = 0.25, n = 2170 phases including 1135 onsets). Together, these results support the hypothesis (**Figure 3B**) that slow, rhythmic fluctuations in arousal interact with reward history to determine the timing of exploration onset.

### Pupil size predicts flattened neural tuning in prefrontal cortex during exploration

To probe the neural mechanisms linking pupil size to exploration, we examined how pupil size predicts neural activity in the FEF (Bruce and Goldberg, 1985; Moore and Armstrong, 2003; Moore and Fallah, 2001; Schall and Hanes, 1993) (**Figure 4A**). We previously reported that exploration is associated with flattened tuning for choice in FEF neurons (Ebitz et al., 2018). While FEF neurons often predict upcoming choices during exploitation, many show reduced choice selectivity during exploration. Pupil size predicted similar changes in in FEF neurons and did so beyond what could be explained by exploratory states themselves. Out of 155 recorded single neurons, 88 (57%) were tuned for choice (**Figure 4B**; 57%, one sample proportion test: p < 0.001). These are referred to as “tuned neurons,” regardless of whether they were modulated by pupil size. Among tuned neurons, 21 (24%) were also modulated by pupil size, and 16 (18%) showed a significant interaction between choice and pupil size. On average, tuning curves flattened as pupil size increased in both tuned and untuned neurons (**Figure 4C-D**). Among untuned neurons, an additional 22% (15/67) were significantly modulated by pupil size (*p* < 0.05), with a median regression coefficient (β) of –0.0011 ± 0.063. This suggests that pupil-linked mechanisms affect FEF activity even in neurons that are not directly involved in encoding choice. This may suggest a more domain-general role for arousal in modulating prefrontal network dynamics.

**Figure 4.**
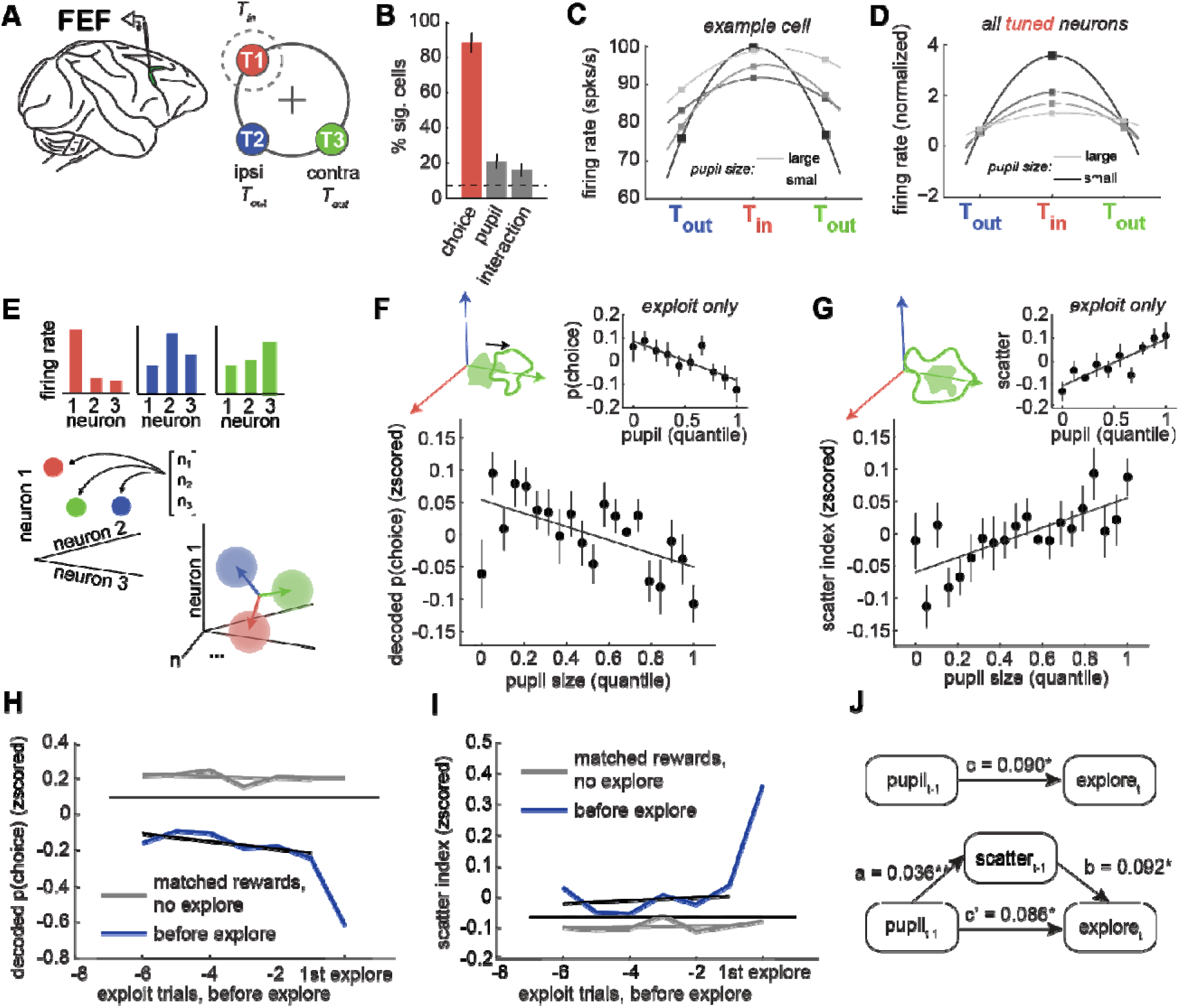
Pupil size predicts choice tuning curves and population disorganization. A) Recordings were made in the FEF. Right: The cartoon illustrates the relative positions of the receptive field target (T_in_, red) and the ipsilateral and contralateral targets (T_out_, blue and green). B) Percent of neurons with significant tuning for choice target, pupil size, and the interaction. C) Tuning curve for an example neuron across target locations, separated by pupil size. Lighter = larger pupil. The dashed line indicates the number of neurons expected to be significant by chance at p < 0.05. D) Same for all tuned neurons, which refers to the 88 out of 155 FEF neurons that were significantly tuned for choice, regardless of their modulation by pupil size. E) Cartoon illustrating how neural population measures consider patterns of firing rates across neurons as vectors in neural state space. Targeted dimensionality reduction is used to find the hyperplane where the distribution of neural activity across trials best predicts choice. Vectors here are the coding dimensions that separate the choices. F) The decoded choice probability (projection onto the correct coding dimension) plotted as a function of pupil size quantile. Inset: Same for exploit trials alone. G) The scatter index, a measure of the variance in choice-predictive population activity, plotted as a function of pupil size quantile. Inset: Same for exploit trials. H) Decoded choice probability for trials before the onset of exploration (in blue) and trials with matched rewards (in gray).I) Scatter index for trials before the onset of exploration (in blue) and trials with matched rewards (in gray). Error bars and shaded regions ± SEM. J) Mediation analysis between pupil size, scatter index, and the onset of exploration. Top: Direct model. Bottom: Indirect, mediated model. Asterisks marked significant paths (*p < 0.01 **p < 0.001).

Because single neurons are noisy, it is difficult to dissociate the effects of pupil size and exploration at the level of individual cells. However, changes in tuning at the single-neuron level also imply shifts in the organization of the neural population and looking at the population level can allow us to estimate these effects within smaller subsets of the data (Ebitz and Hayden, 2021); **Figure 4E**). Indeed, we found that pupil size also predicted changes in how accurately choice information could be decoded from simultaneously recorded populations of FEF neurons. Consistent with our prior results (Ebitz et al., 2018), decoded choice probability was significantly lower during exploration compared to exploitation (paired t-test: both subjects, p < 0.0001, **Figure S2A**). Critically, larger pupil size predicted weaker choice encoding both across all trials (**Figure 4F**; GLM: beta = -0.032, p < 0.0001). This was not driven by differences between the states because pupil size also predicted choice decoding accuracy within exploit trials alone (GLM: beta = -0.037, p < 0.005). There was no significant effect of pupil size on choice decoding within explore trials (GLM: β = –0.003, p = 0.77, **Figure S2B**), which could have been due to differences in trial counts (i.e., explore trials made up only ∼15% of total trials) or to floor effects (decoding accuracy was already close to chance in these trials). These findings show that pupil-linked arousal predicts the strength of choice-predictive neural signals in FEF above and beyond what can be explained by differences between the states.

In our previous work, we found that decreases in choice-predictive activity were accompanied by increases in variability in FEF population responses to the same choice (Ebitz et al., 2018). We quantified this with the “scatter index”: a measure of the spread within clusters of same-choice population activity (see **Methods**). A high scatter index indicates that neural activity on a given trial was dissimilar to other trials where the same choice was made, whereas a low scatter index indicates that neural activity was tightly clustered. We observed a higher scatter index during exploration compared to exploitation (paired t-test: both subjects, p < 0.0001, **Figure S2C**). Here, we also found that increasing pupil size predicted an increase in the scatter index in both subject B and subject O (**Figure 4G**; GLM: beta = 0.04, p < 0.0001). This effect remained significant and of similar magnitude within exploit trials alone (GLM: β = 0.03, *p* < 0.0005), again suggesting that the relationship between pupil size and scatter was not an artifact of state differences with pupil size. Pupil size again did not significantly predict the scatter index during explore trials (GLM: β = 0.0006, *p* = 0.94, **Figure S2D**). Thus, pupil size predicted disorganization of choice-predictive signals in the FEF, at both the level of single neurons and in the population.

### Neural disorganization mediates the relationship between pupil size and exploration

To test whether neural population activity, like pupil size, also specifically predicted the onset of exploration, we compared its dynamics in the trials preceding exploration to those in matched-reward control trials. While sudden changes in the decoded choice probability and scatter index were largely aligned with the onset of exploration (as reported previously), these neural measures were at a different average level on the trials preceding exploration, compared to reward-matched controls (**Figure 4H-I**; choice probability, offset = -0.611, p < 0.001, n = 28; scatter index = 0.258, p < 0.001). Reward information did not cause a change in either variable (choice probability, slope = -0.004, p > 0.5; scatter index = 0.002, p > 0.5), while small, but significant interaction terms suggested that both variables anticipated the onset of exploration (choice probability interaction = -0.058, p < 0.01; scatter index interaction: beta = 0.040, p < 0.001). To determine if these neural measures might explain or mediate some of the relationship between pupil size and exploration, we turned to structural equation modeling (Preacher et al., 2007; Sobel, 1986). We found that the scatter index was a significant mediator of the relationship between pupil size and the onset of exploration (**Figure 4J**; effect of mediation, ab = 0.003, p < 0.005; full report in Table S1). Together, these results suggest that pupil size predicts disruptions in the organization of prefrontal neural activity that then mediate its relationship with the onset of exploration.

### Exploration may reflect a critical transition in brain state dynamics

Neural systems, like other complex networks, can undergo tipping points—irreversible “critical transitions” between stable operating regimes (O’Byrne and Jerbi, 2022; Scheffer, 2020; Scheffer et al., 2009; Wang et al., 2012). Because exploration occurs as the brain passes from exploiting one target to exploiting another, it is worth considering the possibility that exploration may represent a critical transition in brain states. Indeed, during exploration, we previously reported (Ebitz et al., 2018) several phenomena in the FEF and in behavior that are hallmarks of critical transitions, including a rapid flickering back and forth between choices (Wang et al., 2012), an increase in the variance in neural activity (Scheffer et al., 2009), and a disruption of long-term neuronal autocorrelations that suggests that passing through exploration causes time-irreversible changes in the FEF network (Scheffer, 2020). However, there is another classic feature of critical transitions that we did not consider: an early warning signal known as “critical slowing”. As the system nears the tipping point, the dynamics within the system begin to flatten out in preparation for the change. As a result, the systems’ processes slow down and take longer to trace the same paths (Scheffer et al., 2009). Therefore, we next asked if there was any evidence that decision-making slowed down in advance exploration in this dataset.

To test for critical slowing, we examined two measures of decision speed: one behavioral and one neural. First, we looked at response time, a measure of how long it takes the brain to generate saccadic decisions. Response time was not only slower in the trials before exploration, compared to matched-reward control trials (**Figure 5A-C**; GLM offset = 0.39, p < 0.0001, n = 28), but it slowed down over trials before the onset of exploration (interaction = 0.05, p < 0.001). Second, we looked at the mean rate of change in neural population choice signals during the decision process (“neural speed”, see **Methods**). Neural speed was only weakly correlated with response time across sessions (mean = -0.07, min = -0.36, max = 0.09, Pearson’s correlation), suggesting that the measures were complementary, rather than redundant. Like response time, neural speed was also significantly slower on average in the trials before exploration, compared to matched-reward controls (**Figure 5D–F**; GLM offset = -0.17, p < 0.0001, n = 28). However, unlike response time, neural speed did not show a significant slowing trend over trials (interaction = -0.01, p = 0.08). Although the notion that the brain may be subject to critical tipping points is controversial (O’Byrne and Jerbi, 2022), these results are consistent with the idea that exploration could reflect a critical transition between exploiting one option and exploiting another.

**Figure 5.**
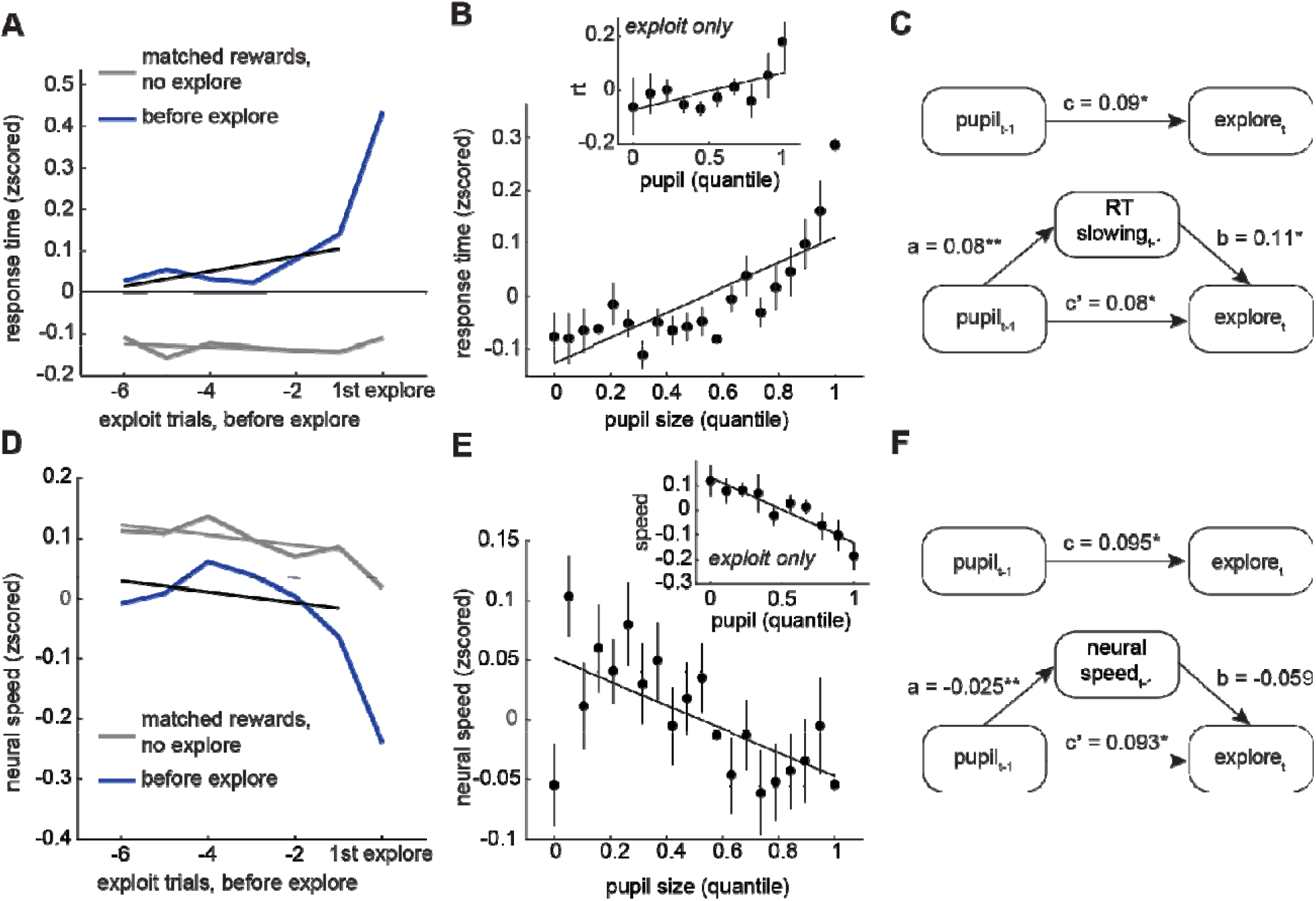
Pupil size predicts behavioral and neural slowing. A) Response time on exploit trials before the onset of exploration (blue) and trials with matched rewards but no exploration (gray). B) Response time plotted as a function of pupil size quantile. Inset: Same for exploit trials alone. C) Mediation analysis between pupil size, response time, and the onset of exploration. Top: Direct model. Asterisks marked significant paths (*p < 0.01). Bottom: Indirect, mediated model. D) Neural speed on exploit trials before the onset of exploration (in blue) and trials with matched rewards (in gray). E-F) Same as B-C for neural speed. Shaded areas and error bars ± SEM.

We first asked whether slowing effects could be better explained by the typical reward histories that precede exploration, rather than internal states like arousal. However, reward history alone did not have a significant effect on either neural or behavioral slowing (response time: slope of matched-reward trials = 0.0002, p > 0.5; neural speed: slope = -0.018, p > 0.1). This suggests that some internal variable, like arousal, could be driving increased slowing and, perhaps, also the systems’ proximity to a tipping point. Indeed, increasing pupil size predicted slower response times (**Figure 5B**; GLM beta = 0.08, p < 0.0001, n = 28 sessions), even within periods of exploitation (beta = 0.05, p < 0.0001). The same was true of neural slowing (**Figure 5E**; all trials: beta = -0.03, p < 0.0005; exploit only: beta = -0.09, p < 0.0001). Further, structural equation modeling revealed that both measures of slowing mediated the relationship between pupil size and the onset of exploration (**Figure 5C and F**; Table S2-3). In sum, the pupil-linked mechanisms that anticipated exploration included both a disorganization of neural activity and a slowing of decision-related computations in brain and behavior—hallmarks of a system approaching a critical transition.

## Discussion

Random decision-making is a powerful strategy for exploration (Dayan and Daw, 2008; Ebitz et al., 2018; Gershman, 2019; Wilson et al., 2021, 2014) that is linked to disorganized patterns of neural activity in the prefrontal cortex (Ebitz et al., 2018; Muller et al., 2019; Wilson et al., 2021). Here, we sought to identify some of the neurobiological mechanisms that drive random exploration and its neural signatures. We found that pupil size, a peripheral correlate of autonomic arousal, predicted exploration and certain measures of neural population activity previously linked to exploration. Consistent with previous studies (Jepma and Nieuwenhuis, 2011a), pupil size was generally larger during exploration, compared to exploitation. However, there was also a complex temporal relationship, where pupil size ramped up between periods of exploration and decreased during exploration. As a result, pupil size was largest at the beginning or “onset” of exploration and explained variance in the onset of exploration that could not be explained by other variables. Together, these results suggest that pupil-linked mechanisms may play a role in driving the brain into an exploratory state.

Our behavioral results largely replicate previous findings linking exploration to increased pupil size (Jepma and Nieuwenhuis, 2011a). However, where we found gradual ramping before exploration and sudden constriction after, Jepma and Nieuwenhuis (2011) reported an abrupt (if modest) increase of pupil size at the onset of exploration and then a gradual decrease at the return to exploitation. The discrepancy may be due to differences in the operational definition of exploration. Jepma and Nieuwenhuis (2011) fit an RL model to behavior and defined “explore choices” as the choices that were not reward-maximizing according to the model. This definition conflates exploration with errors of reward maximization. A strategy that is non-reward-maximizing would produce choices that are orthogonal to value, not consistently bad. Here, we used an HMM to identify latent explore and exploit states on the basis of the temporal profiles of choices alone, with no assumptions on the underlying value computations. This allowed us to dissociate the effects of reward history from the explore/exploit choice labels. We reported here (**Figure 1C**), and in previous studies (Chen et al., 2021; Ebitz et al., 2018), that HMM labels outperform RL labels in explaining behavioral and neural measures, suggesting that the HMM may more accurately separate distinct neural and behavioral states. If the HMM allows for more precise identification of exploratory and exploitative choices, it would follow that it also allows for more precise reconstruction of the temporal relationship between the pupil and exploration.

The precision of our explore/exploit labels revealed that the U-shaped relationship between pupil size and exploration was caused by a refractory constriction in the pupil. When exploration was plotted as a function of pupil size, the relationship appeared non-linear: both small- and large-pupil choices were more likely to be exploratory. This superficially resonated with the idea of a U-shaped relationship between arousal and task performance (i.e. the “Yerkes-Dodson curve”; (Aston-Jones and Cohen, 2005; Yerkes and Dodson, 1908): perhaps reliable exploitation is only possible at intermediate levels of arousal. However, when we examined the temporal relationship between exploration and pupil size, we found that pupil size only predicted the onset of exploration, the first explore choice in a sequence. Small-pupil explore choices happened because starting to explore seemed to “reset” the level of pupil-linked arousal, causing it to quickly fall below baseline. If increased pupil size promotes a transition to exploration, then it is possible that post-exploration constriction represents a refractory period for exploration. Given that uncertainty grows with time in this task (and in all dynamic environments), it may not be smart to start to explore again immediately after you have just explored. A refractory period could ensure that non-reward-maximizing explore choices are deployed only when needed. Future work is needed to test this hypothesis and to determine the cognitive and/or neurobiological mechanisms at play.

Before exploration, we observed an oscillatory dynamic that was about twice as fast as the 10 trials it took the pupil to recover after exploration. This 0.06-0.09 Hz oscillation entrained the onset of exploration: onsets tended to occur during the rising phase of pupil size, whereas identical trial sequences that did not result in exploration were on the opposite phase. This implies that it is the confluence of pupil size, pupil phase, and trial history that best predicts the onset of exploration. This result reinforces the idea that arousal or arousal-linked mechanisms help trigger random exploration (Ebitz and Moore, 2019; Gilzenrat et al., 2010; Reimer et al., 2016), rather than just tracking the reward-linked variables that make exploration more probable. It is also notable that the period of the pupil oscillation was close to the frequency of the Mayer wave: an oscillation in blood pressure that entrains other autonomic measures, including respiration and heart rate (Borjon et al., 2016a; Japundzic et al., 1990; Julien, 2020, 2006; Kamiya et al., 2005). There is precedent for the idea that behavior can be entrained by the Mayer wave: in marmosets, fluctuations in arousal predict the spontaneous onset of a call (Borjon et al., 2016a). This paper argued that the Mayer wave may function to organize vocal communication by bringing the system closer to the threshold for transitioning from inaction to action. It is possible that oscillations in the pupil and pupil-linked mechanisms function the same way here, organizing important state changes in time. In parallel, pupil-linked mechanisms seem to anticipate other state transitions, including belief updating (Filipowicz et al., 2020; O’Reilly et al., 2013), task disengagement (Kane et al., 2017), and other behavioral state changes (Bouret and Sara, 2005). Together, these results suggest an important role for pupil-linked mechanisms in driving successful transitions between certain neural and behavioral states.

Critically, pupil size and pupil oscillations did not predict all state transitions here, but only the transition into exploration. What kinds of state transitions might be entrained by pupil-linked arousal? It is possible that the pupil may have a special relationship with certain “critical” kinds of transitions. Critical transitions are abrupt, large-scale, and irreversible changes in the dynamics and behavior of complex systems, like the brain. As these systems go from being in one conformation (i.e. always choosing the left option) into another conformation (i.e. always choosing the right), the system dynamics that support the old state have to disappear and the new dynamics have to emerge. During this brief transitory period, when both dynamics co-exist in the system, certain signatures can be observed in the system. We previously reported that the exploration was accompanied by abrupt changes in neural population activity, certain patterns of noise in brain and behavior, and disruptions in long-term neuronal autocorrelations: all observations that could be interpreted as suggesting that exploration is a critical transition in the brain (Ebitz et al., 2018). Here, we found that pupil size predicts these features of neural activity and also an prominent “early warning sign” of critical transitions: a slowing, in brain and behavior, of the decision process. While there are certain patterns of activity in FEF that predict response speed (Hauser et al., 2018; Yao and Vanduffel, 2023), here we identified *independent* neural and behavioral measures of decision speed that both mediated the relationship between pupil size and exploration. Notably, pupil size also predicted slower neural and behavioral responses within exploit-only trials, suggesting that these effects are not an artifact of differences between explore and exploit states. This suggests that these effects may reflect a domain-general influence of arousal on cognitive dynamics, consistent with the idea that fluctuations in pupil-linked neuromodulation shape the temporal structure of decision-making in general, beyond any role in state transitions. In this view, the transition into exploration in FEF may reflect an extremum of these domain-general arousal effects—a tipping point—rather than signals that are highly specific to state transitions. Together these results suggest that pupil-linked arousal pushes neural and behavioral states to a critical tipping point and highlights the crucial role of pupil-linked mechanisms in changing the dynamics of the brain.

What underlying, pupil-linked mechanisms could support critical transitions? Changes in pupil diameter coincide with neuromodulator system activity, especially norepinephrine (NE) and acetylcholine (Breton-Provencher and Sur, 2019; de Gee et al., 2020; Joshi et al., 2016; Joshi and Gold, 2020; Murphy et al., 2014; Reimer et al., 2016). At the neuronal level, central NE flattens tuning curves, at least in the auditory cortex (Martins and Froemke, 2015), though it may have different effects in non-cortical structures (Manella et al., 2017). Here, we made a parallel observation: as pupil size increases, neuronal turning curves_flattened and choice-predictive neural population activity became disorganized. These results resonate with a particularly influential theory of NE function: the idea that NE release may facilitate “resets” in cortical networks in order to effect long-lasting changes in brain and behavior (Aston-Jones and Cohen, 2005; Bouret and Sara, 2005). More recent studies seem to consistently report that elevated levels of NE predict an increase in behavioral variability, while pharmacological blockade of NE receptors reduces variability (Chen et al., 2023; Kane et al., 2017; Sadacca et al., 2017; Tervo et al., 2014). In combination with the present study, these results could suggest that phasic NE signaling functions to push the brain towards a critical tipping point where it is better able to transition from one regime to another. In this view, behavioral variability would be linked to NE not because NE increases variability directly, but because the brain is more likely to transition into a high variability regime after it is released. Of course, pupil size is also associated with other neuromodulatory systems, cognitive factors, and other measures of arousal. Thus, future work is needed to identify the neurobiological mechanisms that underpin the relationship between pupil size and critical transitions that we report here.

## Materials and Methods

### Surgical and electrophysiological procedures

All procedures were approved by the Stanford University Institutional Animal Care and Use Committee. Subjects were two male rhesus macaques, surgically-prepared with head restraint prostheses, craniotomies, and recording chambers under isoflurane anesthesia via techniques described previously (Ebitz et al., 2018). Following surgery, analgesics were used to minimize discomfort, and antibiotics were delivered prophylactically. After recovery, subjects were acclimated to the laboratory and head restraint, then placed on controlled access to fluids and trained to perform the task.

In order to train the animals on the explore/exploit task a gradual procedure was used in which the two animals were first trained to make saccadic eye movements in exchange for liquid rewards. Once the animals reliably made controlled eye movements to a single target (generally within 1–2 days), a second target was introduced, and the animals were free to choose between them. At the outset, each target was associated with a probability of reward (initially 10% and 90%), which was reversed in blocks at the experimenter’s discretion. Over a period of 2–4 months, the difference in reward probabilities between the targets was gradually reduced, the blocks transitioned into gradual reward probability shifts (reward walks), and a third target was introduced. The speed and order of these changes depended on each animal’s performance and engagement with the task. One animal (monkey O) was naïve to laboratory tasks prior to this experiment, whereas the second (monkey B) had been previously trained on covert and overt attention tasks, but not on any prior value-based tasks.

Recording sites were located within the FEF, which was identified via a combination of anatomical and functional criteria. The location of recording sites in the anterior bank of the arcuate sulcus was verified histologically in one subject and via microstimulation in both subjects (Ebitz et al., 2018). Recordings were conducted with 16-channel U-probes (Plexon), located such that each contact was within gray matter at an FEF site. An average of 20 units were recorded in each session (131 single units, 443 multi units; 576 total units across 28 sessions).

### General behavioral procedures

Eye position and pupil size were monitored at 1000 Hz via an infrared eye tracking system (SR Research; Eyelink). The manufacturer’s standard methods for calculating pupil area were used. MATLAB (Psychtoolbox-3; (Kleiner et al., 2007)) was used to display stimuli and record behavioral responses and pupil size measurements. Task stimuli were presented against a dark gray background (7 cd/m2) on a 47.5 cm wide LCD monitor (Samsung; 120 Hz refresh rate, 1680 x 1050 resolution), located 34 cm in front of the subject.

### Three-armed bandit task

The subjects performed a sequential decision-making task in which they chose between 3 targets whose values changed over time. The subject first fixated a central fixation square (0.5° stimulus, +/- 1.5-2° of error) for a variable interval (450-750ms). At any point within 2s after the onset of the targets, subjects indicated their choice by making a saccade to one of the targets and fixating it (+/- 3°) for 150 ms. Reward magnitude was fixed within session (0.2-0.4 μL). Reward probability was determined by the current reward probability of the chosen target, which changed independently over trials for each of the three targets. On every correct trial, each target had a 10% chance of the reward parameter changing either up or down by a fixed step of 0.1, bounded at 0.1 and 0.9. Because rewards were variable, independent, and probabilistic, the subjects could only infer the values of the different targets by sampling them and integrating noisy experienced rewards over multiple trials.

### General analysis procedures

Data were analyzed with custom software in MATLAB. Unless otherwise noted, all t-tests were paired, two-sided t-tests, and generalized linear models were run on raw data, with session number coded as a dummy variable to account for session-to-session variability. Model comparison was based on standard methods that involve calculating the likelihood of the data and Akaike information criteria (AIC) of each model, then using AIC weights to identify (1) the model that is most likely to minimize information loss, and (2) the relative likelihood of competing models to do the same (Burnham and Anderson, 2004). For analyses of any behavioral or neural variables on the trials before or after exploration, continuous runs of exploit trials were required. The values of behavioral and neural variables were *z*-scored within a session to facilitate comparisons across sessions. In the results section, we refer to a z-score of 0 as “baseline”. The 200 ms window immediately preceding target onset was chosen as the analysis epoch for all choice-predictive neural measures. A longer, whole-trial epoch was chosen for neural speed analyses (0 to 500 ms) following target presentation. Firing rates were computed per trial.

### Pupil size

Pupil size was measured during the first 200 ms of fixation, a time at which the eye was fixed at a known point on the screen, illumination was identical across trials, and anticipatory changes in the pupil were minimal. To remove any blinks or movement artifacts, trials where pupil size or the change in pupil size from the first time bin of this epoch to the last was +/- 6 standard deviations from average were eliminated from further analyses. A total of 178 trials (out of 21,793, approximately 0.8% of observations) were outliers.

### Hidden Markov Model

To identify when subjects were exploring versus exploiting, we employed a hidden Markov model (Chen et al., 2021; Ebitz et al., 2018). In this framework, choices Y_t_ are treated as emissions from a latent decision-making state z_t_, which can either be an explore or one of the multiple exploit states.

The emission model for exploration assumed a uniform probability of selecting any option:

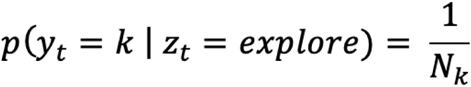

Where N_k_ is the total number of options. In contrast, exploit states deterministically emitted choices to the exploited option i:

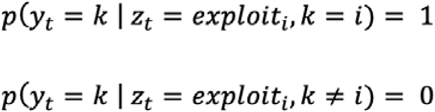

Latent state transitions followed a Markov process, such that the probability of the current state depended only on the previous state:

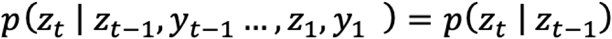

To reduce model complexity, parameters were shared across exploit states, and subjects were assumed to begin in the explore state. The final HMM included only two free parameters: the probability of persisting in exploration and the probability of persisting in exploitation. The model was fit using expectation-maximization with 20 random restarts, and the solution maximizing the observed data log-likelihood was selected. The most probable sequence of latent states was recovered using the Viterbi algorithm.

### Reinforcement learning model

To compare goal state labels derived from an RL and HMM model, we employed a Rescorla-Wagner model. This was fit using maximum likelihood estimation. The value of each option is iteratively updated according to:

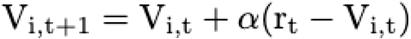

Where *V_i,t_* is the value of option *i* at time *t*, *r_t_* is the reward at time *t*, and α represents the fitted learning rate, which determines how much the difference between the predicted and actual reward (the prediction error) influences value. To make a decision, the values are passed through a softmax decision rule:

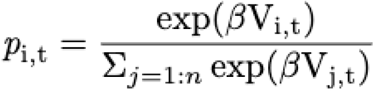

Where *n* is the total number of available options, and β is the inverse temperature, which controls the level of random noise in decision-making. After (Daw et al., 2006; Jepma and Nieuwenhuis, 2011a; Pearson et al., 2009), decisions that were not reward maximizing were labeled as exploratory (i.e. any decision where V_chosen,t_ was not the maximum V at time *t*).

### Generalized Linear Model (GLM)

To examine the relationship between behavioral and neural variables, we employed generalized linear models (GLMs). These models were fit using maximum likelihood estimation. Each dependent variable (i.e., scatter index, response time, or choice probability) was modeled as a linear combination of predictors:

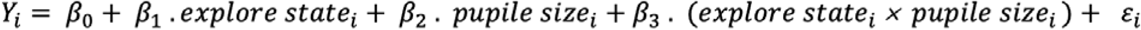

Where Y_i_ is the dependent variable on trial i, β_0_ is the intercept, and β_1_, β_2_, β_3_ are regression coefficients quantifying the influence of the corresponding predictors (e.g., explore state, pupil size, and their interactions).ε_i_ is the residual error term.

In analyses where categorical variables (e.g., explore state: 0 = exploit, 1 = explore) were used as predictors, these were coded as binary dummy variables. Models were fit using the identity link function and assumed normally distributed errors.

### Learning Index

To investigate whether learning differed with pupil size within the exploratory choices, we calculated a learning index that captured the effect of rewards experienced during exploration on future choices. Because reward effects decay exponentially quickly (Lau and Glimcher, 2008), a 1-trial-ahead index should capture most of the variability in how much is learned between trial types. The equation was:

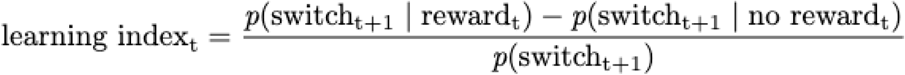

### Lagged change in pupil size

To determine whether exploration impacted pupil size, we measured the change in pupil size (Δ pupil) between pairs of trials that either were or were not separated by at least 1 explore trial. Segments of twenty-five consecutive trials were identified that either included a single bout of exploration or did not include exploration. For each pair of trials within these sequences, we then measured the change in pupil size between the first exploit trial of the sequence (t1) and the remaining exploit trials in the sequence (t2:25). This was repeated for all unique pairs of trials that met our selection criteria.

### Matched reward trials

To test whether the rising trend in pupil size before exploration is best explained by reward history, we identified trial sequences with identical reward and state histories that did not end in exploration (“matched rewards”). For each onset of exploration preceded by at least 6 exploit trials, we searched for identical sequences of exploit trials, with identical reward histories, that did not end in exploration. We chose 6 previous trials because this was the longest sequence of reward history we could regularly match within the majority of sessions (we could find at least 10 matched sequences in 96% [27/28] of sessions for 6 trials sequences; that dropped to 75% [21/28] at 7 trials). Identical results were obtained with other sequence lengths, though these analyses included fewer sessions.

### Mediation analysis

To determine if the predictive relationship between pupil size and exploration was mediated by other variables, we used structural equation modeling to test for mediation. Mediation analyses involve fitting three regression models. The first model measures the total effect (c) of the independent variable (here, pupil size) on the independent variable (here, onset of exploration):

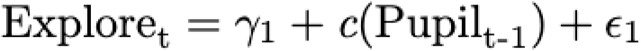

In these equations, represents the intercept for each equation, while L represents the error of the model. Note that the estimated parameter c will include both direct effects of pupil size on exploration, but also indirect effects that may be mediated by other variables. Therefore, we also fit a second model that tests if the independent variable also predicts a potential mediator variable (here, neural network scatter):

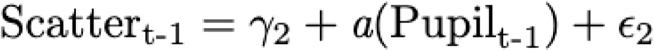

Model parameter a thus captures the effect of pupil size on the mediator. Finally, a third model estimates the unique contributions of both the potential mediator (scatter, b) and the independent variable (pupil size, c’), now controlling for the mediator:

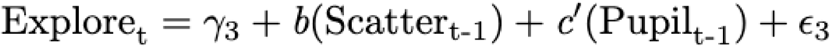

A drop between c and c’ indicates that the effect of the independent variable (pupil) on the dependent variable (exploration) is reduced when the mediating variable is considered. The mediation effect (the indirect effect of the pupil size on the onset of exploration via the mediating factor) can also be estimated directly, via taking the product of the coefficients a and b. Sobel’s test is used to determine the significance of the mediation path (Sobel, 1986).

### Phase analysis

To determine if the onset of exploration happened at a specific phase of pupil size over trials, we performed a wavelet analysis. Because this method only assumes local stationarity, it is more suitable than other methods for analyzing pupil size, which tended to ramp over trials. A wavelet was constructed by multiplying a complex sine wave (frequency = 5 trials) with a Gaussian envelope (μ = 0, σ = cycles / (2π*frequency), cycles = 5; (Cohen, 2014)). The wavelet was convolved with the baseline pupil size time series and the phase of the signal was calculated on each trial (Matlab; angle). Standard circular statistics were used to measure the differences between phase distributions for explore onsets and reward-matched controls (Zar, 1999) and the phase alignment within these trial types (Berens, 2009). The latter was also verified via comparison with bootstrapped null distributions (1000 samples).

### Targeted dimensionality reduction

Neural state spaces have as many dimensions as there are recorded neurons, but converging evidence suggests (1) that the neural states that are observed in practice are generally confined to a lower-dimensional “manifold”, and (2) that task-relevant information is encoded by a small number of dimensions in that manifold. Because we wanted to isolate the effects of arousal on choice-related activity from well-known effects of arousal on neural activity (Ebitz and Platt, 2015; McGinley et al., 2015; Pfeffer et al., 2022; Podvalny et al., 2021; Reimer et al., 2016, 2014; van Kempen et al., 2019; Waschke et al., 2019), we focused all our neural population analyses on activity within the choice-predictive subspace, rather than on neural activity more broadly.

To do this, we used targeted dimensionality reduction to identify the choice-predictive dimensions of the neural state space (Cohen and Maunsell, 2010; Cunningham and Yu, 2014; Ebitz et al., 2018; Peixoto et al., 2021). Specifically, we used multinomial logistic regression (Matlab; mnrfit, mnrval, (Hastie et al., 2009)) to identify the separating hyperplanes that best discriminated each choice from the alternative choices. This is equivalent to fitting a system of binary classifiers of the form:

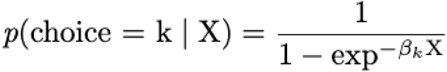

Where one classifier discriminates target 1 choices from targets 2 and 3 and a second discriminates target 2 choices from targets 1 and 3. The classifier that discriminates target 3 from targets 1 and 2 is then just the negative of target 1 and target 2. These axes span the subspace in which neural activity best predicts choice. Classifiers were trained on firing rates from an epoch that began when the targets appeared and ended at the time of the saccade. Mean imputation was used for the small number of occasions where a unit was not held for the whole duration of the session (∼3% of trials, ∼12% of units) and a small fraction of units were omitted from these analyses because their mean firing rates were less than 2 spikes/s, which makes their weights difficult to identify (∼8% of units).

### Choice Probability Decoding

Within the choice-predictive subspace, the distance from the separating hyperplanes (the vectors illustrated in **Figure 4E**) are the decoding vectors: the vectors along which we can project neural activity in order to decode the log odds of choice. This projection is equivalent to the decoded choice probability from the multinomial logistic regression model and this is the figure we took as the decoded choice probability in **Figures 3F** and **3H**. We evaluated decoding accuracy by measuring how often the most-likely choice predicted by the model coincided with the choice the subject made.

### Scatter index

The scatter index measures how much choice-predictive population neural activity is clustered between trials with the same choice (Ebitz et al., 2018). It is calculated by measuring the average Euclidean distance of each trial from all other trials where the same choice was made and dividing it by the average Euclidean distance to all other trials where a different choice was made:

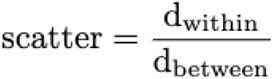

Each trial thus has its own scatter index value, with a value of 1 indicating no difference in clustering between same-choice and different-choice trials, and a value less than 1 indicating greater clustering with same-choice trials compared to different-choice trials.

### Neural speed

To determine how the speed of the decision-making process changed before and during exploration, we calculated the rate of change in neural states within the choice-predictive subspace during the first 400 ms following target presentation. Each trial’s neural activity was sampled in non-overlapping 20 ms bins and then projected into the choice-predictive subspace. The change in neural activity within the subspace was then calculated between each pair of samples. Finally, the changes were averaged together across the trial and normalized to the bin width to produce an average rate of change in choice-predictive activity for that trial.

## Acknowledgments

This work was supported by the Natural Sciences and Engineering Research Council of Canada (Discovery Grant RGPIN-2020-05577), the Fonds de Recherche du Québec–Santé (Junior 1 Chercheur-Boursier 284309 to R.B.E.), the Jacobs Foundation (Research Fellowship, seed grant to R.B.E.), and CIFAR Azrieli Global Scholars (seed grant to R.B.E.), the National Eye Institute (R01-EY014924 to T.M.), and l’Institut de valorisation des données (fellowship to K.J.).

---

**Table S1.** Regression coefficients and p values for the mediation analysis testing whether the scatter index mediates the relationship between pupil size and the onset of exploration. Related to **Figure 4J**.

|  |  | pupil<sub>t-1</sub> → scatter<sub>t-1</sub> → explore<sub>t</sub> |  |
| --- | --- | --- | --- |
|  |  | est. coefficient | p value < |
| Total effect | c | 0.090 | 0.005 |
| Effect on mediator | a | 0.036 | 0.0005 |
| Unique mediator effect | b | 0.092 | 0.005 |
| Indirect effect | ab | 0.003 (z = 2.70*) | 0.005 |
| Direct effect | c' | 0.086 | 0.005 |

\*Sobel's test

**Table S2.** Regression coefficients and p values for the mediation analysis testing whether response time slowing mediates the relationship between pupil size and the onset of exploration on the next trial. Related to **Figure 5C**.

|  |  | pupil<sub>t-1</sub> → RT slowing<sub>t-1</sub> → explore<sub>t</sub> |  |
| --- | --- | --- | --- |
|  |  | est. coefficient | p value < |
| Total effect | c | 0.090 | 0.005 |
| Effect on mediator | a | 0.078 | 0.0001 |
| Unique mediator effect | b | 0.106 | 0.0005 |
| Indirect effect | ab | 0.008 (z = 3.48*) | 0.0005 |
| Direct effect | c' | 0.080 | 0.01 |

\*Sobel's test

**Table S3.** Regression coefficients and p values for the mediation analysis testing whether neural slowing mediates the relationship between pupil size and the onset of exploration on the next trial. Related to **Figure 5F**.

|  |  | pupil<sub>t-1</sub> → neural slowing<sub>t-1</sub> → explore<sub>t</sub> |  |
| --- | --- | --- | --- |
|  |  | est. coefficient | p value < |
| Total effect | c | 0.095 | 0.005 |
| Effect on mediator | a | -0.025 | 0.0005 |
| Unique mediator effect | b | -0.059 | 0.06 |
| Indirect effect | ab | 0.001 (z = 1.69*) | 0.05 |
| Direct effect | c' | 0.093 | 0.005 |

\*Sobel's test

**Figure S1.**
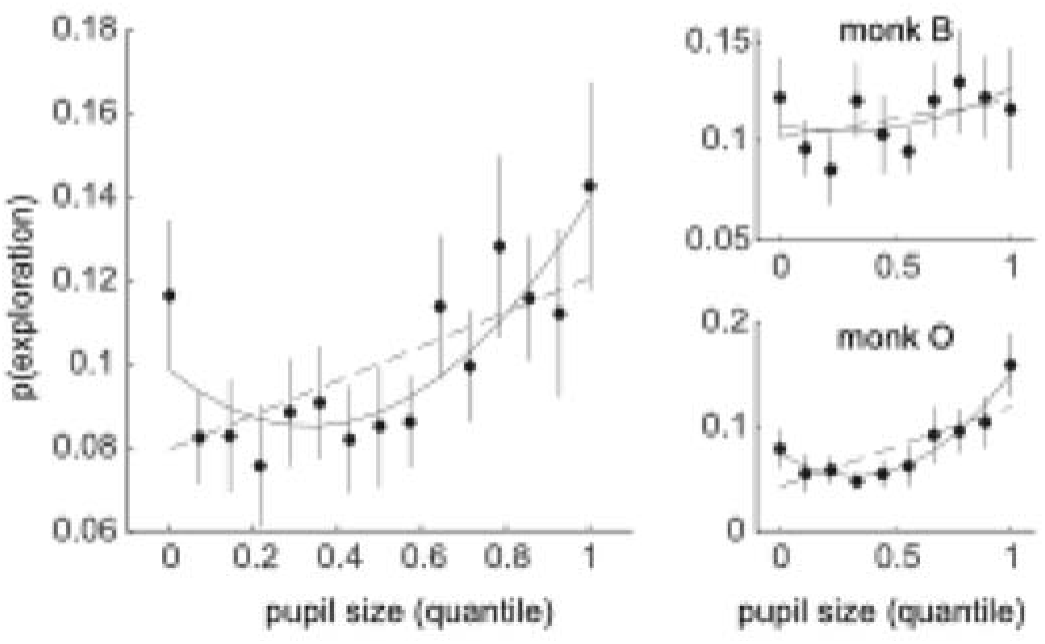
Same as Figure 1E, but without first explore trials.

**Figure S2.**
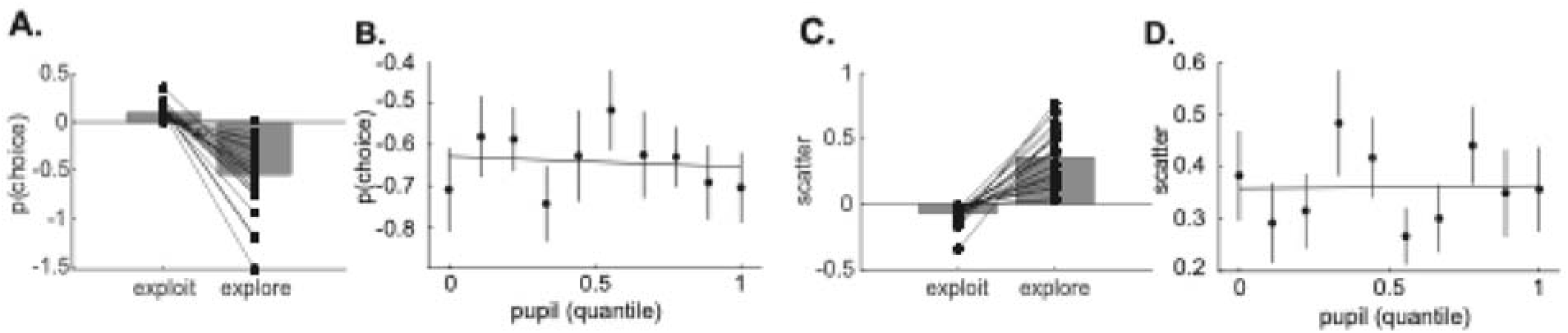
Decoded choice probability and scatter index across behavioral states and pupil size. (A) Decoded choice probability (projection onto the correct coding dimension) for exploit and explore states. Dots represent individual sessions, with lines connecting values from the same session across states. (B) Decoded choice probability plotted as a function of pupil size quantile for explore trials alone. (C) Scatter index, a measure of variance in choice-predictive population activity, for exploit and explore states, with lines connecting values from the same session across states. (D) The scatter index plotted as a function of pupil size quantile for explore trials.

